# AI reveals a damage signalling hierarchy that coordinates different cell behaviours driving wound re-epithelialisation

**DOI:** 10.1101/2024.04.10.588842

**Authors:** Jake Turley, Francesca Robertson, Isaac V. Chenchiah, Tanniemola B. Liverpool, Helen Weavers, Paul Martin

## Abstract

One of the key tissue movements driving closure of a wound is re-epithelialisation. Earlier wound healing studies have described the dynamic cell behaviours that contribute to wound re-epithelialisation, including cell division, cell shape changes and cell migration, as well as the signals that might regulate these cell behaviours. Here, we use a series of deep learning tools to quantify the contributions of each of these cell behaviours from movies of repairing wounds in the *Drosophila* pupal wing epithelium. We test how each is altered following knockdown of the conserved wound repair signals, Ca^2+^ and JNK, as well as ablation of macrophages which supply growth factor signals believed to orchestrate aspects of the repair process. Our genetic perturbation experiments provide quantifiable insights regarding how these wound signals impact cell behaviours. We find that Ca^2+^ signalling is a master regulator required for all contributing cell behaviours; JNK signalling primarily drives cell shape changes and divisions, whereas signals from macrophages regulate largely cell migration and proliferation. Our studies show AI to be a valuable tool for unravelling complex signalling hierarchies underlying tissue repair.

## Introduction

Tissue damage triggers a complex series of overlapping cell and tissue movements that recapitulate embryonic morphogenetic episodes, which together will stave off infection and eventually repair the wound back to something approaching its pre-wounded state (Gurtner et al., 2008; Eming et al., 2014; Peña & Martin, 2024). One of the key tissue movements contributing to wound healing is re-epithelialisation, whereby the epidermal cells at the cut wound edge and the sheet of epithelium behind them advance to seal the wound gap. Individual and concerted cell contractions and shape changes, as well as cell movements/migration and cell divisions, all contribute to the closure of the epithelial gap (Abreu-Blanco et al., 2012; Park et al., 2017; Aragona et al., 2017; Razzell et al., 2014; Tetley et al., 2019; Turley et al., 2023) but how much each of these cell behaviours contributes to the ultimate goal is not known; neither do we know precisely which signals of those known to be activated soon after tissue damage (e.g., the wound calcium wave, JNK signalling, and signals released by wound inflammatory cells), might regulate each cell behaviour, nor how any of these cell behaviours might be compensated for if one or more of the others fail.

In recent years, there have been several elegant live imaging studies of wound re-epithelialisation in mouse models using sophisticated state-of-the-art imaging strategies to enable the capture of cell movements and cell divisions within the advancing epidermis. One of these studies targeted wounds in the murine tail, where the epidermis is devoid of distracting hairs (Aragona et al., 2017). This study provided definitive evidence that the front rows of wound edge cells migrate whilst those further back proliferate (Aragona et al., 2017). Another study, this time utilising the thin hairless epidermis of the ear, showed the first high-resolution imaging of an advancing wound epidermis. This too indicated zones of migration at the leading edge and proliferation further back, and this data was of sufficiently high resolution to enable tracking of fluorescently-labelled epidermal nuclei and quantify both migratory speed and rates of proliferation (Park et al., 2017).

Because tissues in mouse and human are opaque, any live imaging approaches are, by definition, technically extremely challenging. Before the studies described above, it had not been feasible to directly observe mammalian re-epithelialisation, and even these pioneering new studies provide rather limited volumes of data and offer less than optimal spatial and temporal resolution. As a complement to murine approaches, several recent studies have utilised the translucency and genetic tractability of *Drosophila* embryos, larvae, and pupae to investigate wound re-epithelialisation (Razzell et al., 2014; Tetley et al., 2019; Tsai et al., 2018; Turley et al., 2023; Wu et al., 2009). These studies have been useful in revealing the contributions not just of leading edge epithelial cells, where a contractile actomyosin purse-string helps draw these cells forward, but also of cells back from the leading edge where tissue fluidity - driven by cells jostling within the sheet - is pivotal for releasing tissue tension within the “following” epithelial sheet, and thus enabling the front row cells to advance forward (Razzell et al., 2014; Tetley et al., 2019).

Collecting large movie datasets of healing wounds is more achievable in translucent models such as *Drosophila* and zebrafish but requires new methodologies for analysis of the data (Turley et al., 2022). Indeed, recently, we and others have used deep learning algorithms (U-NetCellDivision) to precisely quantify when and where cells are dividing within unwounded tissues and in the advancing wound epidermis within the *Drosophila* pupa (Turley et al., 2023; Villars et al. 2023). We used another deep learning algorithm (U-NetBoundary) to segment the cell boundaries, enabling us to reveal how daughter cells shuffle post division to align with the global tension in the tissue (Turley et al., 2023).

In the present study, we use deep learning models to integrate this cell division data with the two other cell behaviours that contribute to wound re-epithelialisation, cell shape changes and cell motility. We do this by using additional deep learning algorithms to detect cell divisions and the shapes of cells in the wound epithelium; in parallel, we also quantify cell migration by tracking the nuclei of cells in time-lapse movies using a single-particle tracking algorithm (Turley et al., 2023).

Ultimately, we would like to determine the contributions of each cell behaviour to the wound closure effort and how each of these cell behaviours is regulated by conserved wound-induced signals. To that end, here we have analysed large data sets of the three cell behaviours extracted from live time-lapse confocal microscopy of wounded *Drosophila* pupae. We first characterised the typical re-epithelialisation of wounds in wild type tissue, before genetically manipulating several well documented wound healing signals - the calcium wave (the first damage signal released after wounding (Razzell et al., 2013; Sipka et al., 2021; Tu et al., 2018)), JNK signalling (Hammouda et al., 2020; Weavers et al., 2019), and finally the innate immune inflammatory response (Thuma et al., 2018; Weavers et al., 2018; Weavers et al., 2020) - in order to determine which of these sources of signals might mediate the regulation of which of these cell behaviours.

## Results

The wing of the *Drosophila* pupa is an ideal model for investigating the cell biology of wound healing because it is immobile and optically translucent, enabling long term, high-resolution imaging, and also because of its considerable genetic tractability (Etournay et al., 2015; Popović et al., 2017; Weavers & Wood, 2016). 18 APF pupae (APF, after prepuparium formation) were wounded with a laser after removal of the outer pupal case (Fig. 1A). We generate these lesions in a consistent region of the wing (Fig. 1B) because mechanical stresses inevitably will vary across the developing wing and can influence wound re-epithelialisation (Etournay et al., 2015; Paci & Mao, 2021; Popović et al., 2017; Turley et al., 2023). The wing at this developmental stage consists of two flattened epithelial sheets with intervening hemolymph, in which resides innate immune cells (called hemocytes in *Drosophila*) and fat body cells ((Franz et al., 2018) and Fig. 1C). Wounds were generated in pupae expressing *E-cadherin-GFP* and *Histone2-RFP* (to label epithelial cell boundaries and nuclei, respectively), with a micropoint laser, and subsequently imaged by confocal microscopy with 3D stacks taken every 2 minutes for 3 hours.

**Figure 1.**
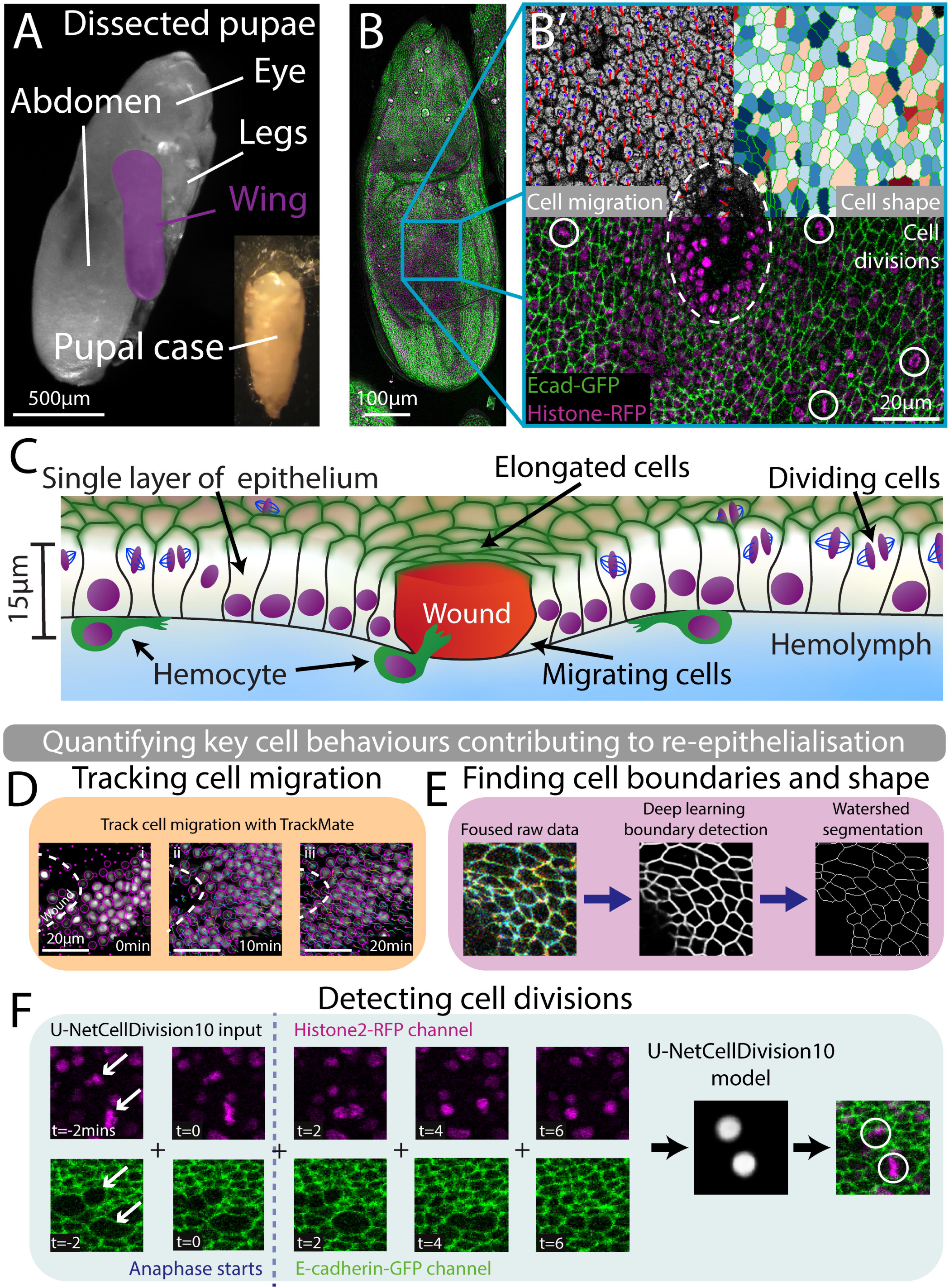
Live imaging of pupal wing wounds and automated quantification of the 3 cell behaviours contributing to re-epithelialisation. A) Bright field image of an 18hour APF (after puparium formation) pupa with the wing highlighted in magenta. B) Tile scan of the full pupal wing. E-cadherin-GFP (green) shows cell boundaries and Histone2-RFP (red) labels cell nuclei. B’) Wounded wing tissue with white dashed line showing the wound edge. The 3 cell behaviours involved in wound healing are shown in 3 segments. Top left (cell migration), shows a close-up of the wounded epithelium in greyscale with cell nuclei marked with a mauve circle and the velocity of each nucleus indicated by a red line. The length of the line is proportional to the speed of that nucleus. In the top right (cell shape), cells elongated perpendicular to the wounds and highlighted in blue and red are cells oriented towards the wound. Bottom (cell divisions) have been detected and highlighted with a white circle. C) Diagram of cross section of wounded 18h APF pupal wing. D) Snap shots from a movie of the wounded epithelium at timepoints indicated. Purple circles highlight detected nuclei and purple dots indicate a detected nuclei above or below the plane of view. White dashed lines indicate the wound edge. Nuclear tracks show cells moving towards the wound. E) A 3-focal plane image is inputted into the U-NetBoundary model and then segmented using Tissue Analyzer F) Our deep learning algorithm for detecting cell divisions in the wound epithelium. The model input has 10 frames, 5 each from the Histone2-RFP and E-cadherin-GFP channels. The model highlights (with a white spot) where it detects a division.

### Automated AI algorithms enable us to quantify each of the three cell behaviours that contribute to wound closure

We tracked and quantified the three cell behaviours that contribute to wound re-epithelialisation – cell migration, cell shape changes and cell divisions – by means of automated algorithms. We used the plugin TrackMate to track the nuclei of cells over time in 3D utilising the *Histone2-RFP* channel data (Tinevez et al., 2017). Cell migration was measured by averaging the radial component of the velocity of a cell’s nucleus at given times after wounding and at different distances from the leading edge (Fig. 1D). Next, we applied the first of our deep learning models, U-NetBoundary (Turley et al., 2023), to detect cell boundaries (from the *E-cadherin-GFP* channel), and thus identify individual cells (Fig. 1E). Cell elongation relative to the wound was calculated by computing individual q-tensors for each cell from their cell boundaries (Olenik et al., 2023). These q-tensors comprise a component for cell elongation and another for a cell’s orientation (see Methods for further details). A heatmap of cell elongation relative to the wound indicates which cells are elongated perpendicular to (pointing towards) the wound edge in red and those aligned along the wound margin appear blue (Fig. 1B’). To quantify the shape of cells adjacent to the wound, we averaged the cells’ q-tensors, binning them into groups of cells as a function of distance from the wound, and the time after wounding.

Similarly, we detected cell divisions within the wounded epithelium with a deep learning model called U-NetCellDivision (Turley et al., 2023), which utilises dynamic cell boundary and nuclear information from the GFP and RFP channels (Fig. 1F). We binned these dividing cells into radial bands (annuli) extending out from the wound edge, and for various time points post wounding, to determine division density for each tissue annulus back from the wound edge.

### Epithelial cells extending several cell diameters back from the leading edge contribute to wound closure

To determine how wound size impacts the cell biology of repair, we generated wounds of two different sizes; large wounds had an initial area between 700-1100μm^2^, and small wounds were between 200-400μm^2^ (Turley et al., 2023). The large wounds took, on average, 50mins to close to 20% of their initial size, and small wounds healed the same amount in 25mins (Fig. 2A). At this late stage of wound closure, both wound sizes have leading edge cells that make contact with one another to seal the wound closed (Abreu-Blanco et al., 2012; Wood et al., 2002). This last part of wound closure is visually very noisy, and the leading edge becomes unclear. Hence, we define a wound as closed at this point as the majority of re-epithelialisation has occurred (see Fig.2B-B’’ for time course of a large wound healing and Supplemental Movies S1 and S2 for repair of small and large wounds, respectively).

**Figure 2.**
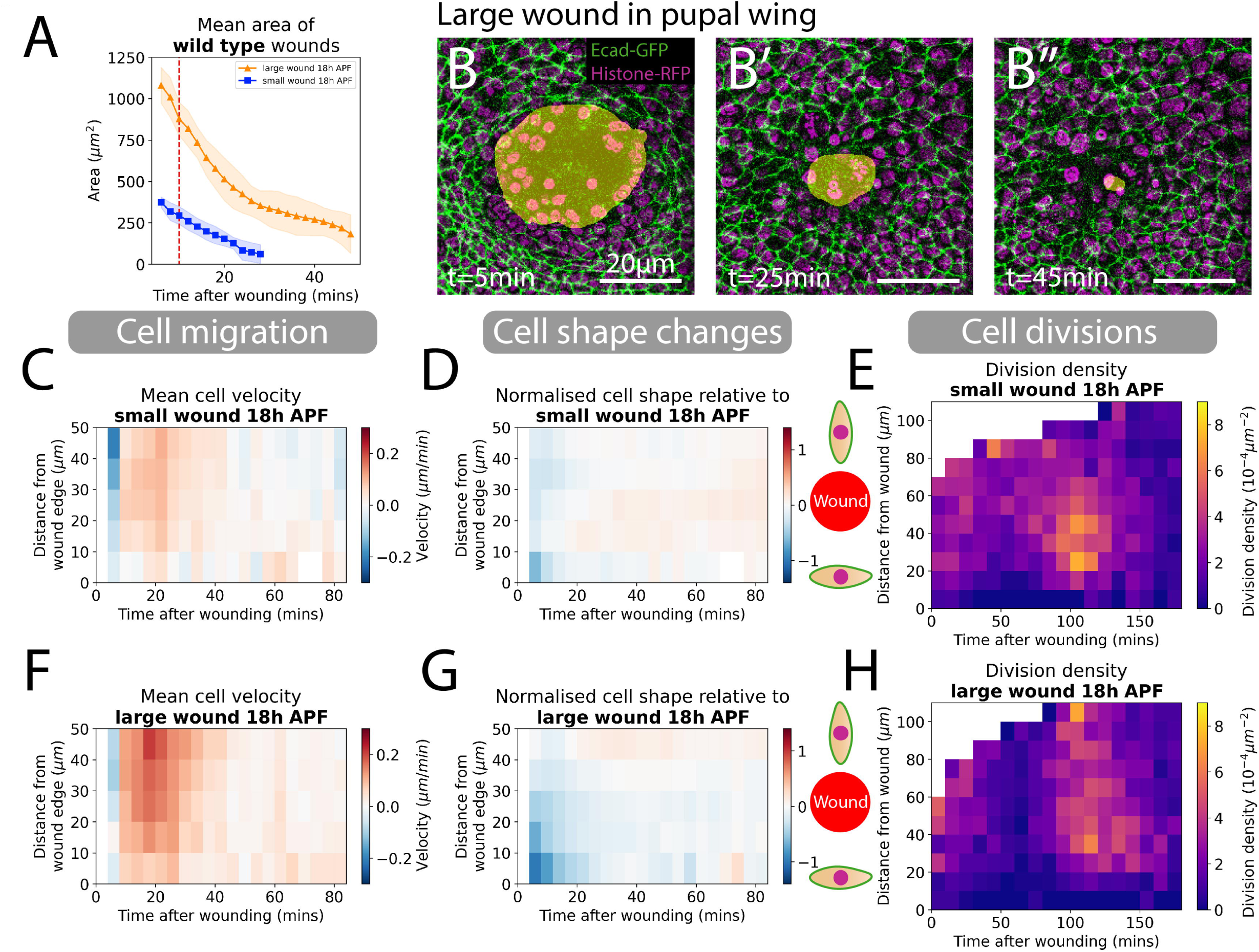
Quantifying cell migration, shape changes and divisions around small and large wounds. A) Quantification of the wound area over time for both small and large wounds. The red dashed line is the 9-10 minute point where we classify the size of wounds. B-B”) A large wound as visualised over 3 timepoints 5mins, 25mins, and 45mins. Yellow shading highlights non-epithelial tissue in images to illustrate wound closure over time. C,F) Spatio-temporal heatmaps of cell velocity relative to wound centre for small and large wounds, respectively. Blue regions indicate cells are migrating away from wounds and red cells are migrating towards the wound. D,G) Spatio-temporal heatmaps of cell shape elongation relative to wounds for small and large wounds, respectively. Blue regions indicate that cells are elongated perpendicular to wounds and red cells are oriented towards the wound. E,H) Heatmaps of the division density for small and large wounds. (n=8 small wounds and n=9 large wounds)

We next quantified the spatiotemporal distribution of each of the three contributing cell behaviours during wound closure. Our heatmaps indicate the mean velocity of cell nuclei towards the wound as a function of distance from the wound edge and time after wounding, with red highlighting cells migrating towards the wound and blue away from it (Fig. 2C, F). For small wounds, cell rows extending up to 50μm (or 10+ rows of cells), back from the leading edge, migrate toward the wound; this migration persists throughout the 25 mins of healing time. For larger wounds, cells migrate faster and for a longer time, which is as expected due to the longer distance needed to be travelled for effective closure of these wounds (Fig. 2C, F). Interestingly, there are two distinct phases of migration observed during closure of large wounds; for the first (approximately 30 mins post-wounding) cells migrate rapidly, but subsequently migration slows over the next ≈15 minutes as the wound closes (Fig. 2C, F).

Next, we examined the cell shape changes during wound closure. In the cell shape heatmaps we calculated the mean q-tensor of the cells for each distance from wound and time after wounding. This was used to determine cell elongation relative to the wound, with cells elongating towards the wound centre indicated by red in the heatmap, and those elongating along the wound edge highlighted in blue (Fig. 2D, G, and see Methods for details). The q-tensor is a dimensionless quantity, and we have normalised its value using the maximum absolute value of cell elongation around large wounds. For small wounds, cells at the leading edge, and one row back from this edge, rapidly elongated perpendicular to the wound margin (Fig. 2D); these cells subsequently became rounder and were no longer elongated after 10 minutes; this shape reversion contributes to wound closure. In large wounds too, cell elongation occured in leading edge cells after wounding. But here, cell elongation propagated back much further into the neighbouring tissue, with cells up to 30μm (approx. 7-8 rows) back from the leading edge affected (Fig. 2G). Similar to small wounds, cells rapidly reverted back to a rounder shape such that by 25 mins post wounding the mean cell shape had returned to the unwounded tissue average (Fig. 2G). This initial period of cell shape change aligns with the early ‘fast phase’ of cell migration (Fig. 2F).

### A synchronised surge in cell divisions occurs after wounding in a zone back from the leading edge

The third cell behaviour contributing to wound re-epithelialisation is cell division. We quantified the density of proliferation around wounds as a function of space and time, and this data is presented as heatmaps in (Fig. 2E, H and (Turley et al., 2023)). During the initial period, when cells are rapidly migrating and changing their shapes, we see a clear suppression of cell division in the wound edge epithelium, reaching a minimum at 60mins. This suggests that cell proliferation and the other contributing wound cell behaviours might be mutually exclusive, which is also suggested in studies of mammalian wounds (Aragona et al., 2017; Park et al., 2017). For small wounds, the zone of suppressed cell divisions extends 30μm back from the wound edge and lasts until 80 minutes after wounding (Turley et al., 2023). In larger wounds, this suppression of cell divisions has a similar time course, but extends over a much larger area, stretching back over 100μm (>25 cell rows) back from the wound, beyond the field of view. After these periods of reduced cell division, we observe a synchronised burst of proliferation beginning 90 minutes after wounding and lasting for an hour (Turley et al., 2023). This occurs in an annulus ranging from 20 to 70μm back from the small wound sites and 20 to >100μm for large wounds (Fig. 2E, H). This surge of proliferation could be a consequence of synchronised wound signals inducing cells to divide to replenish cells lost during wounding, and/or cells that would have divided if not restricted by wounding becoming synchronised as suppression ends.

### A rapid calcium wave travels back through the wounded epithelium that is required for optimal wound-induced cell migration, cell shape changes and cell divisions

After characterising re-epithelialisation of wounds in wild-type pupal wing tissue, we generated transgenic flies in which key wound healing signals had been perturbed. We began by blocking the well-characterised calcium wave, which is the first damage signal released after tissue injury (Razzell et al., 2013; Weavers et al., 2019). This was achieved by expressing *trpm^RNAi^* to knock down the Trpm calcium channels through which this signal propagates (Razzell et al., 2013); this effectively inhibits the spread of the wound Ca^2+^ wave (Fig. 3E,E’). As ubiquitous expression of *trpm^RNAi^* is partially lethal we temporally controlled its expression using the *Gal4-UAS* system with a temperature sensitive *Gal80^ts^* (McGuire et al., 2004). We kept embryos and larvae at 18°C during early development (to prohibit RNAi expression before pupal stages) and raised the temperature to 29°C once at the pre-pupal stage to activate *trpm^RNAi^* expression. Since flies develop faster at higher temperatures, pupae were raised for 14.75hours at 29°C (instead of 18hours at 25°C) to achieve an equivalent level of development. To confirm consistency of developmental stage, we used cell elongation as a proxy for wing development (Athilingam et al., 2021; Etournay et al., 2015), and found no significant difference between the temperature shift paradigms (Fig. S1).

**Figure 3.**
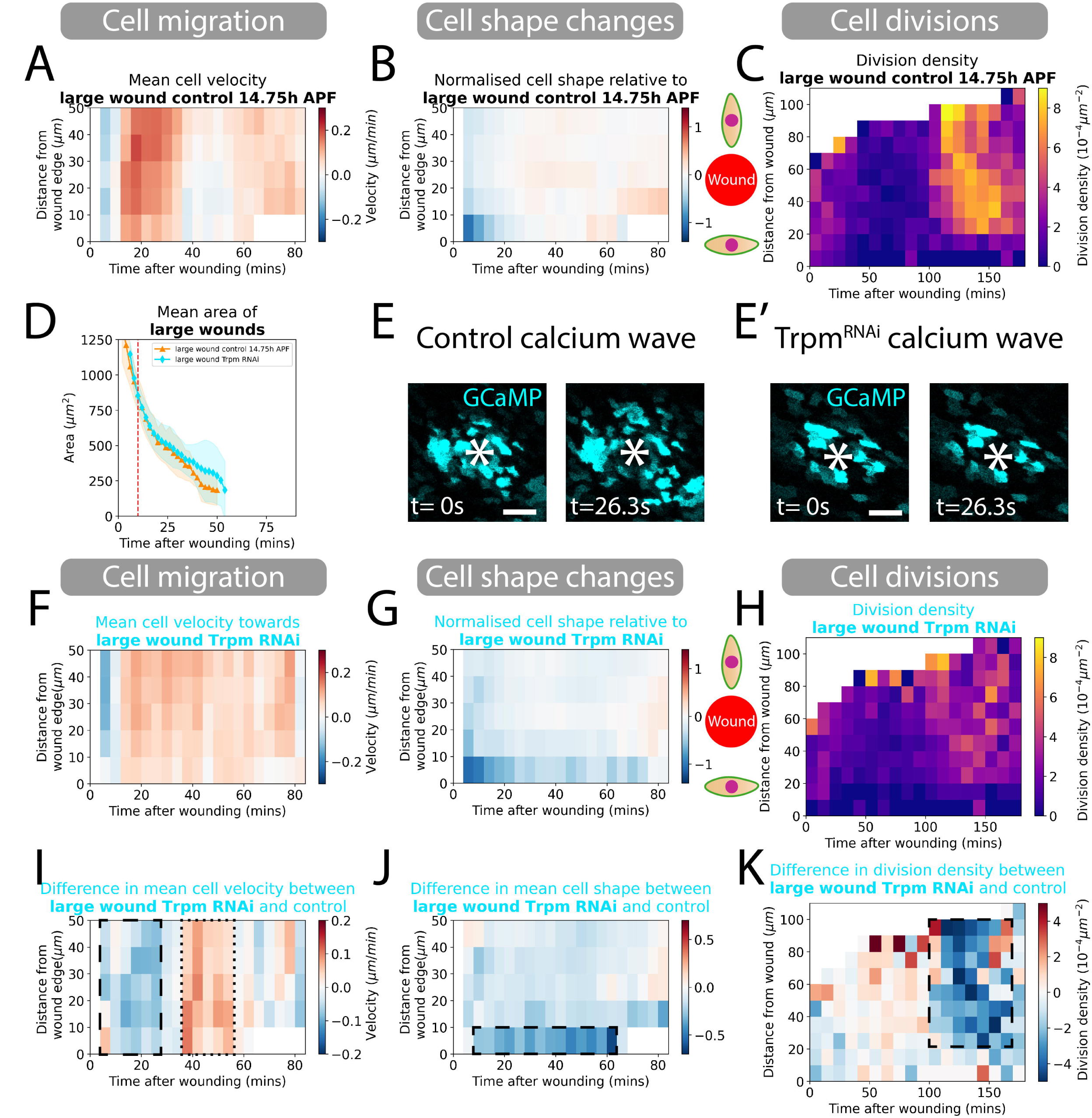
Blocking the wound calcium wave and quantifying the resulting changes in cell behaviours. A) Spatio-temporal heatmaps of cell velocity relative to wounds for control 14.75h APF large wounds. Blue regions indicate those cells that are migrating away from the wound and red those migrating towards the wound. B) Spatio-temporal heatmaps of cell shape elongation relative to wounds for control 14.75h APF large wounds. Blue regions indicate cells elongated perpendicular to the wound and red are cells oriented towards the wound. C) Heatmaps of cell division density for control 14.75h APF large wounds. D) Quantification of the wound area over time as large wounds close. Red dashed line is the 9-10 minute point where we classify the size of wounds. E-E’) Still images from gCAMP movies at 0 and 26 sec post wounding of control versus Trpm^RNAi^ wing tissues to show blocked calcium wave around Trpm^RNAi^ wounds. White asterix indicates location of wound. F) Spatio-temporal heatmaps of cell velocity relative to wound centre for large wounds. Blue regions indicate cells that are migrating away from wounds and red are those migrating towards the wound. G) Spatio-temporal heatmaps of cell shape elongation relative to wounds for large wounds. Blue regions indicate cells that are elongated perpendicular to wounds and red are cells oriented towards the wound. H) Heatmaps of cell division density for large wounds. I) Spatio-temporal heatmaps of the change in cell velocity for control versus wounds with a blocked calcium wave. J) Spatio-temporal heatmaps of the change in cell shape elongation for control versus calcium wave blocked wounds. K) Heatmaps of the change in division density for control versus calcium wave blocked wounds. (n=5 control 14.75h APF wounds, and n=4 Trpm^RNAi^ wounds). Scale bars are 50µm.

We first quantified the area of wounds over time by comparing control (which had been raised with the same temperature shifts) and *trpm^RNAi^* fly lines and observed a slight reduction in the healing rate (Fig. 3D). We went on to investigate whether this gross defect in healing might be linked to any particular changes in the contributing cell behaviour(s). The control (14.75h APF) pupae exhibited a typical wound response with regards to cell migration and shape changes and a clear (delayed) burst of divisions back from the leading edge (Fig. 3A-C). However, all three cell behaviours were altered in *trpm^RNAi^* expressing pupae (Fig. 3F-H). Epithelial cell migration at the wound edge was initially slower for 30 mins after wounding, but migration subsequently persisted for longer than in control wounds, perhaps to compensate for the initially slow start (Fig. 3I). Cell shape changes were also altered, with cells at the leading edge remaining highly elongated for a more extended period (Fig. 3G, J). These two changes in cell behaviour could explain the slower healing of wounds lacking a fully functional calcium wave. We also observe that the synchronised burst of proliferation we observe in wild-type wounds is considerably reduced to approximately half the number of dividing cells in wounded *trpm^RNAi^* pupae (Fig. 3H, K). For all of these genetic perturbation experiments we measured the effect on large wounds since they generally displayed a more extensive cell behavioural response (see earlier), and so we reasoned that any differences after genetic manipulation might be clearer.

### JNK signalling perturbation primarily impacts wound closure by inhibiting changes in cell shape

Next, we disrupted JNK signalling, which is known to be key in developmental morphogenetic episodes such as dorsal closure (Tafesh-Edwards & Eleftherianos, 2020), and in wound healing, both in the formation of actomyosin cables and for associated cell shape changes (Kwon et al., 2010; Lee et al., 2019).

JNK signalling was blocked by expressing a dominant negative mutant version of *Drosophila* JNK (termed *basket, Bsk ^DN^)* in the pupal epithelium as previously utilised for inhibiting JNK signalling (Lee et al., 2019; Tafesh-Edwards & Eleftherianos, 2020; Weavers et al., 2019). We showed that this knockdown was effective by live imaging JNK activity via TRE-DsRed, a transgenic reporter that indicates active JNK signalling, around control and *Bsk^DN^* laser wounds (Fig. 4B,B’). JNK signalling is known to be downstream of the Ca^2+^ wave (Weavers et al., 2019) and so, unsurprisingly, we see a similar, but not identical effect on aspects of wound closure. Again, there was a reduction in overall rate of wound closure with wounds taking 80 minutes to close to 20%, which is half an hour more than for wild type pupal wounds (Fig. (4A)). However, whilst our automated tracking data revealed that rates of cell migration in *Bsk^DN^* pupae were similar to WT tissues (Fig. 4C, F), cells in the wound epithelium appeared unable to effectively change their shape (Fig. 4D, G). Cell elongation around the wound margin failed to propagate back through the tissue. Cells at the leading edge did elongate initially, similar to wild type wounds, but this elongation then took 80 minutes to resolve, rather than only 25 mins in control pupae. This failure of cells to change shape, and thus not contribute to wound closure could be a major reason for the slower healing in pupae with JNK signalling blocked. We also observed a loss of post-wound suppression of cell divisions after JNK knockdown from 30-100mins, as previously described for larval cell death lesions (Cosolo et al., 2019). Our data indicate that while the timing of the wound induced cell proliferative burst is similar to controls the number of dividing cells is reduced (Fig. 4E, H) although to a lesser degree than for the *trpm^RNAi^* pupae.

**Figure 4.**
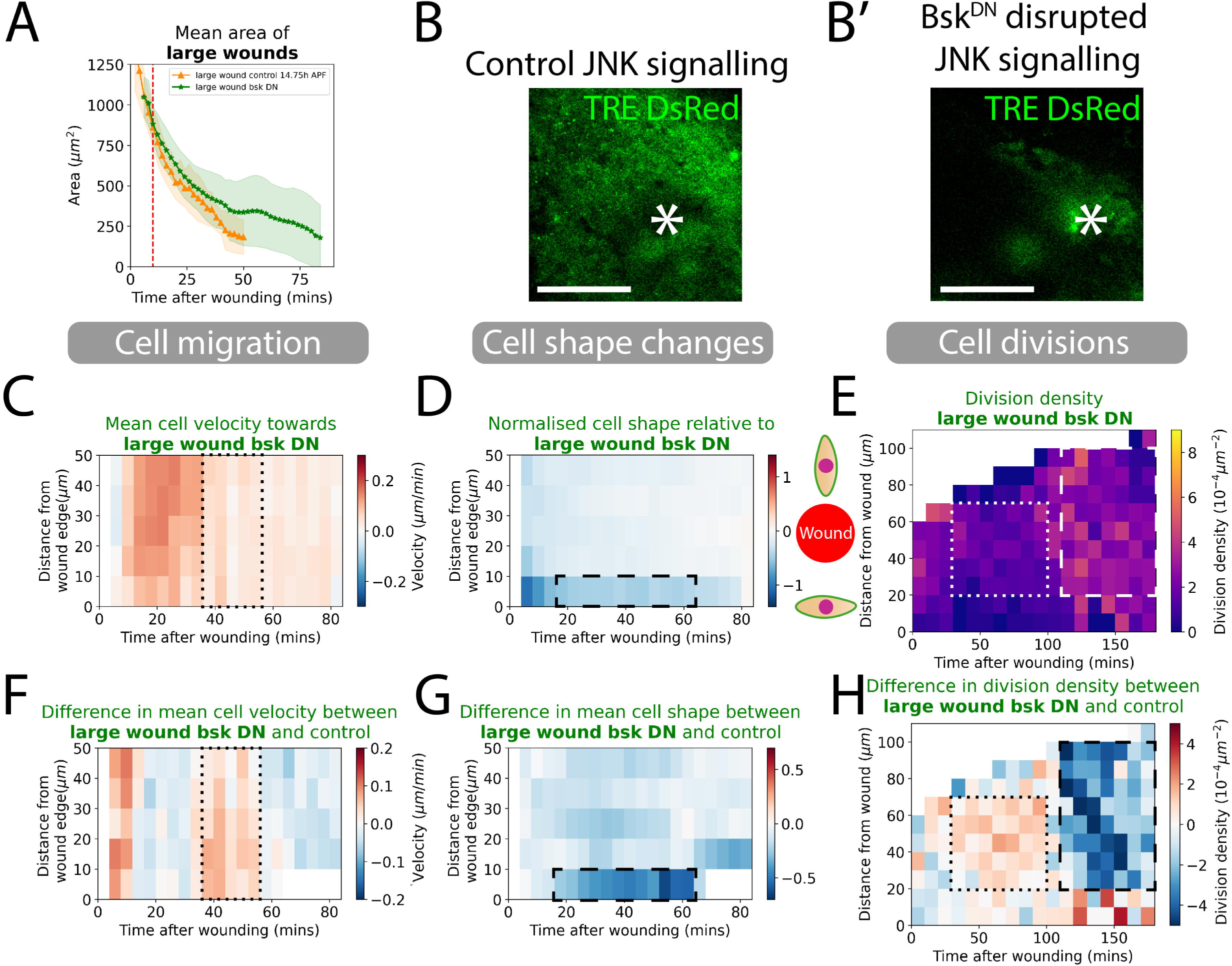
Knockdown of JNK signalling and quantification of how this alters cell behaviours in wound healing. A) Quantification of the wound area over time for control versus JNK knockdown tissues. Red dashed line is the 9-10 minute point where we classify the size of wounds. B-B’) JNK signalling reporter TRE-DsRed in control versus Bsk ^DN^ wing tissues at 5hours post wounding to show dampened JNK signalling around the Bsk ^DN^ wound. White asterix indicates location of wound. C) Spatio-temporal heatmaps of cell velocity relative to wounds for large wounds. Blue regions indicate cells that are migrating away from the wound and red are cells migrating towards the wound. D) Spatio-temporal heatmaps of cell shape elongation relative to wounds for large wounds. Blue regions indicate cells that are elongated perpendicular to wounds and red are cells oriented towards the wound. E) Heatmaps of the division density for wounds. F) Spatio-temporal heatmaps of the change in cell velocity for control versus JNK knockdown wounds. G) Spatio-temporal heatmaps of the change in cell shape elongation for control versus JNK knockdown wounds. H) Heatmaps of the change in cell division density for control versus JNK knockdown wounds. (n=5 control 14.75h APF wounds and n=6 Bsk^DN^ wounds). Scale bars are 50µm.

### Blocking the wound inflammatory response reduces both cell migration and cell proliferation

Lastly, we ablated the flies’ innate immune cells (termed hemocytes) to block the wound inflammatory response (Fig.5). The inflammatory response is believed, through studies in various model organisms, to both clear away wound debris but also release signals that aid in orchestrating various aspects of wound closure including wound re-epithelialisation (Antsiferova & Werner, 2012; Weavers & Martin, 2020). Hemocytes were ablated in pupae by expressing the pro-apoptotic gene, *reaper (rpr)* specifically in hemocytes using *srp-Gal4* in combination with a temperature sensitive Gal80, similar to that performed previously (Weavers et al., 2016); for this genetic manipulation, the shift to 29°C for 14.75hours (to reach an equivalent developmental stage to 18h APF at 25°C) was lethal, and so, instead, we maintained pupae at 18°C for 21hours then shifted them to 29°C for 5hours. Once again, we confirmed there were no statistical differences in developmental stage of control versus “immune ablated” wings (Fig. S1).

**Figure 5.**
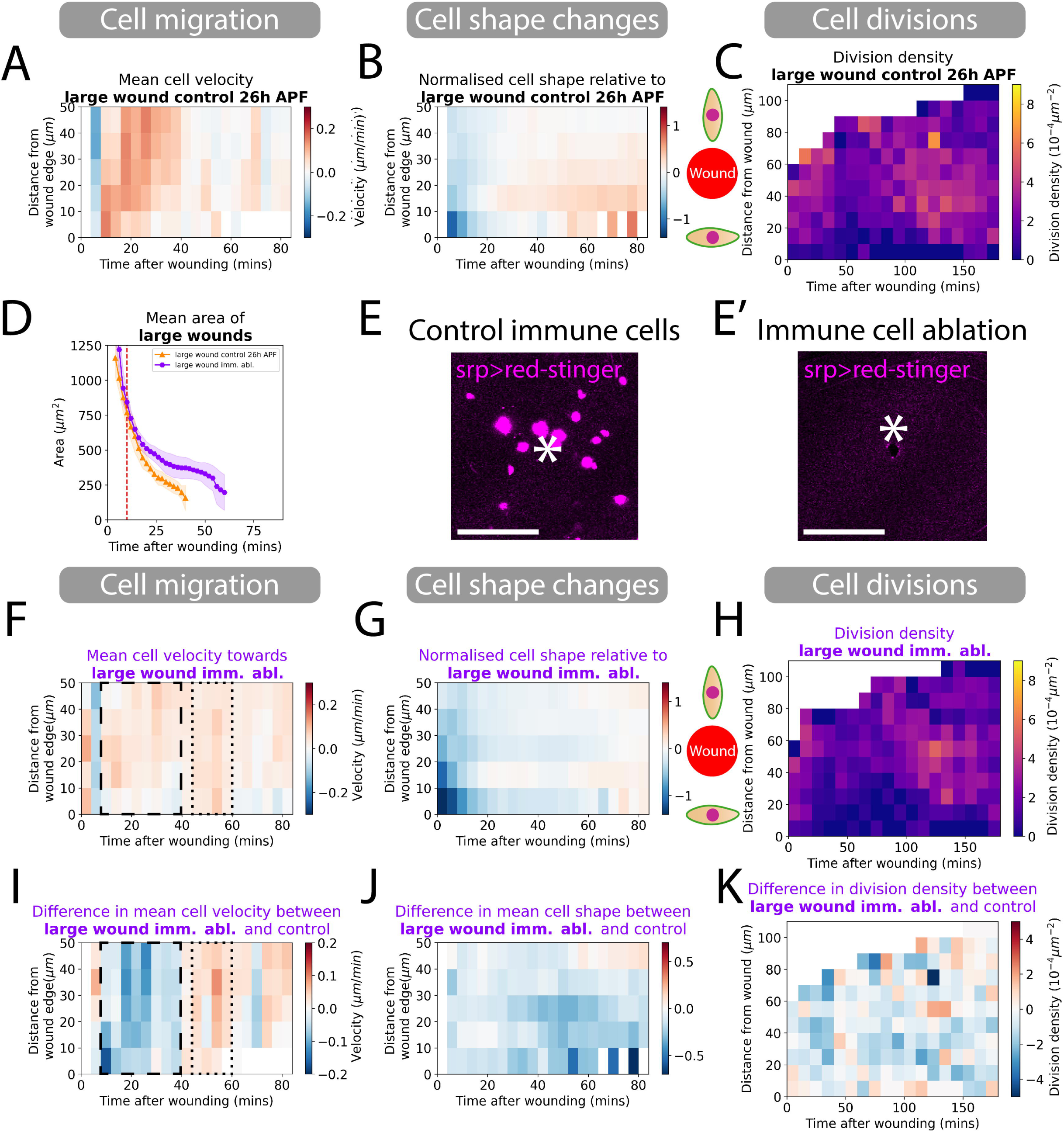
Ablation of immune cells reduces cell migration and decrease divisions globally. A) Spatio-temporal heatmaps of cell velocity relative to wounds for control 26h APF large wounds. Blue regions indicate cells that are migrating away from wounds and red are cells migrating towards the wound. B) Spatio-temporal heatmaps of cell shape elongation relative to wounds for control 26h APF large wounds. Blue regions indicate that cells are elongated perpendicular to wounds and red are cells oriented towards the wound. C) Heatmaps of the division density for control 26h APF large wounds. D) Quantification of the wound area over time for wounds in control versus immune ablated tissues. Red dashed line is the 9-10 minute point where we classify the size of wounds. E-E’) Red-stinger nuclei of innate immune cells recruited to a control wound but absent from immune cell ablated wing tissues. White asterix indicates location of wound. F) Spatio-temporal heatmaps of cell velocity relative to wounds. Blue regions indicate cells that are migrating away from wounds and red cells are migrating towards the wound. G) Spatio-temporal heatmaps of change in cell shape elongation relative to wounds for large wounds. Blue regions indicate cells that are elongated perpendicular to wounds and red are cells oriented towards the wound. H) Heatmaps of the spatial division densities for wounds. I) Spatio-temporal heatmaps of the change in cell velocity for control versus immune ablated wounds. J) Spatio-temporal heatmaps of the change in cell shape elongation for control versus immune cell ablated wounds. K) Heatmaps of the change in cell division density for control versus immune ablated wound. (n=5 control, 26h APF wounds and n=5 rpr wounds). Scale bars are50µm.

We confirmed that this genetic perturbation successfully killed the pupal hemocytes and so prevented the standard immune response to wounds (Fig. 5B, B’). We find that wounds in these immune ablated pupae are also slower to heal, but the contributing disruptions in cell behaviour are subtely different to those where calcium signalling or JNK signalling are blocked (Fig. 5A). This time, cell shape changes remained unaffected, whereas we see that cell migration is severely hindered. We observe almost no cell migration for the first 40 mins and a slow mean nuclei velocity even after 45 mins, by which time control wounds have ceased their cell migration (Fig. 5C, F). In contrast to our studies of repair in JNK knockdown wounds, we observed that cell shape changes are relatively normal, with little difference between control and immune ablation flies (Fig. 5D, G). Wound-induced cell proliferation in these pupae shows the same spatial temporal pattern to that of control wounds but with a clear overall reduction in divisions (Fig. 5E, H). These results indicate that signals from hemocytes appear to be needed for both epithelial cell migration and to stimulate cell division also.

## Discussion

In this study we have developed automated tools to extract information from high-definition videos of epithelial wound closure in the *Drosophila* pupal wing. Our deep learning model takes advantage of both dynamic videos and focused 3D stacks to maximise information input to models for the detection of cell boundaries and nuclei enabling us to quantify cell shape changes, cell migration and cell divisions that together contribute to wound re-epithelialisation. Using these AI methods allows us to generate large datasets and rapidly analyse them with minimal human input.

We quantified the three main epithelial cell behaviours known to contribute to wound healing – cell migration, shape changes and divisions (Aragona et al., 2017; Park et al., 2017) – to determine their contributions to the healing of small and larger wounds. We began by characterising wild-type wound healing. Cell migration and shape changes dominate during the closure of the wounds, while divisions were suppressed initially. Towards the end of healing and after closure, a synchronised burst of divisions occurs, perhaps to regenerate the cells lost during wounding (Turley et al., 2023). Cells migrated quicker and for longer in large wounds compared to small wounds. At the leading wound edge, cells became highly elongated, and this effect is both stronger and propagates further back into the tissue for larger wounds. These cells subsequently changed their shapes, expanding into the wound site to aid in the closure of the wound gap. Only later does cell division resume, with a synchronised burst of proliferation beginning just back from the leading edge and extending further from the wound in larger wounds (Turley et al., 2023).

What damage-induced signals might be regulating these cell behaviours? We have begun to address this systematically in our model, which is amenable to precise quantification of changes in each of these cell behaviours after genetic knockdown of various signalling pathways. We have individually knocked down the first damage signal, the wound calcium wave (Razzell et al., 2013; Weavers et al., 2019), the immediate early JNK response signal which transcriptionally activates many signals downstream of tissue damage (Tafesh-Edwards & Eleftherianos, 2020), and the wound inflammatory response, which is believed to orchestrate many of the tissue responses of wound repair (Antsiferova & Werner, 2012; Weavers & Martin, 2020). Since JNK signalling and macrophage recruitment to wounds are both known to be, at least partially, downstream of calcium signalling we expect to see some degree of overlap in the cell behaviour effects of these knock down studies.

For the first time, we have the tools to test how each of these classes of signals might impact the various contributing wound cell behaviours and indeed compensate for one another. We find that cell migration is slowed when the calcium wave is blocked, and almost stopped in tissues denied a wound inflammatory response, but remains largely unchanged after blocking of JNK signalling (Fig. 4K). Cell shape changes were stalled in wounds where the calcium wave was blocked and almost completely halted in JNK knockdown tissues, but were normal in tissues lacking an inflammatory response (Fig. 6). The synchronised burst of cell divisions in wound epithelium was spatially normal but 50% reduced, in tissues denied a calcium wave. JNK knockdown tissues appeared somewhat desensitised to wounds with regard to cell division, with both a smaller initial suppression and then reduced subsequent burst in divisions; in tissues without an inflammatory response there was a general dampening in proliferation also. Overall, we found that pupal tissues inhibited from experiencing a normal calcium wave displayed a mix of phenotypes reflecting both JNK and inflammation defective flies with regard to their cell migration and shape changes (Fig. 6). This is consistent with the calcium wave, at least in part, being upstream of both of these signals (Razzell et al., 2013; Weavers et al., 2019).

**Figure 6.**
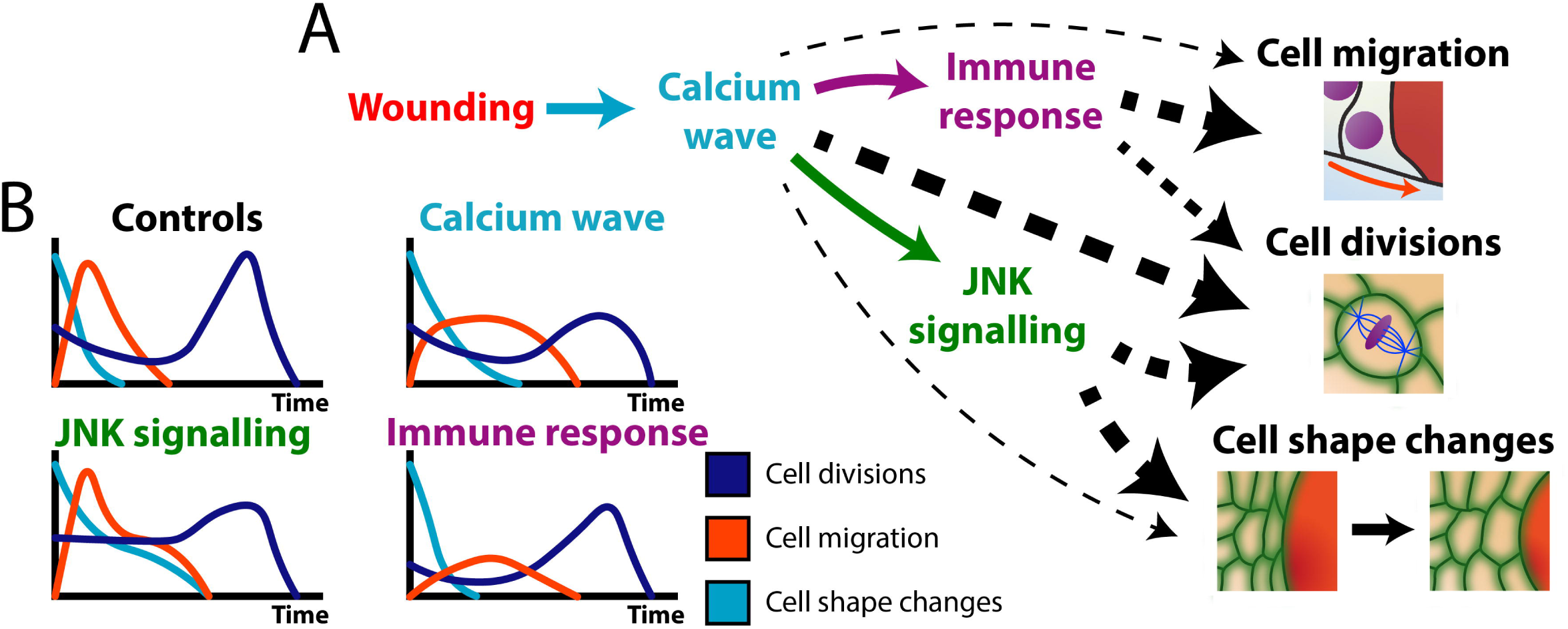
Schematic to illustrate the altered cell behaviours during wound healing in mutant pupal wings in which individual wound signals have been disrupted. A) Schematic of the cascade of wound induced signals and how they impact the cell behaviours that drive re-epithelialisation, as revealed by our study; thickness of dotted back line indicates the relative weighting of the signal. B) Graphical illustrations of the temporal change in cell behaviours during wound healing across wild type and genetic perturbation.

Potential next steps utilising this system will be to examine additional sources of signals and to dissect further with regard to the signals delivered by inflammatory cells, by knocking out individual growth factor/trophic signals from hemocytes, rather than simply ablating the lineage as we have done here. This approach might allow investigations of which pathways the immune cells are using to influence both migration and divisions of epithelial cells. We also could attempt to target each of the cell behaviours one by one, to not only determine their contribution to wound healing but to also study how the other two behaviours are impacted. Can they compensate for the loss of one behaviour, or will they also be negatively impacted?

Applying AI and mathematical tools to biological systems can reveal new results and increase our understanding of underlying mechanisms (Turley et al., 2022). Quantifying properties and behaviours of cells in a tissue on a large scale via automated tools such as deep learning has many applications extending from developmental biology through to cancer. Here, we offer a first insight into what signals regulate each of the cell behaviours that contribute to the evolutionarily conserved process of wound re-epithelialisation. In future, it will be possible to dissect out these signals and how they regulate wound closure in more nuanced ways, and to do similar for the significantly more complex repair of mammalian skin wounds.

## Supplemental Figure Legends

**Figure S1.**
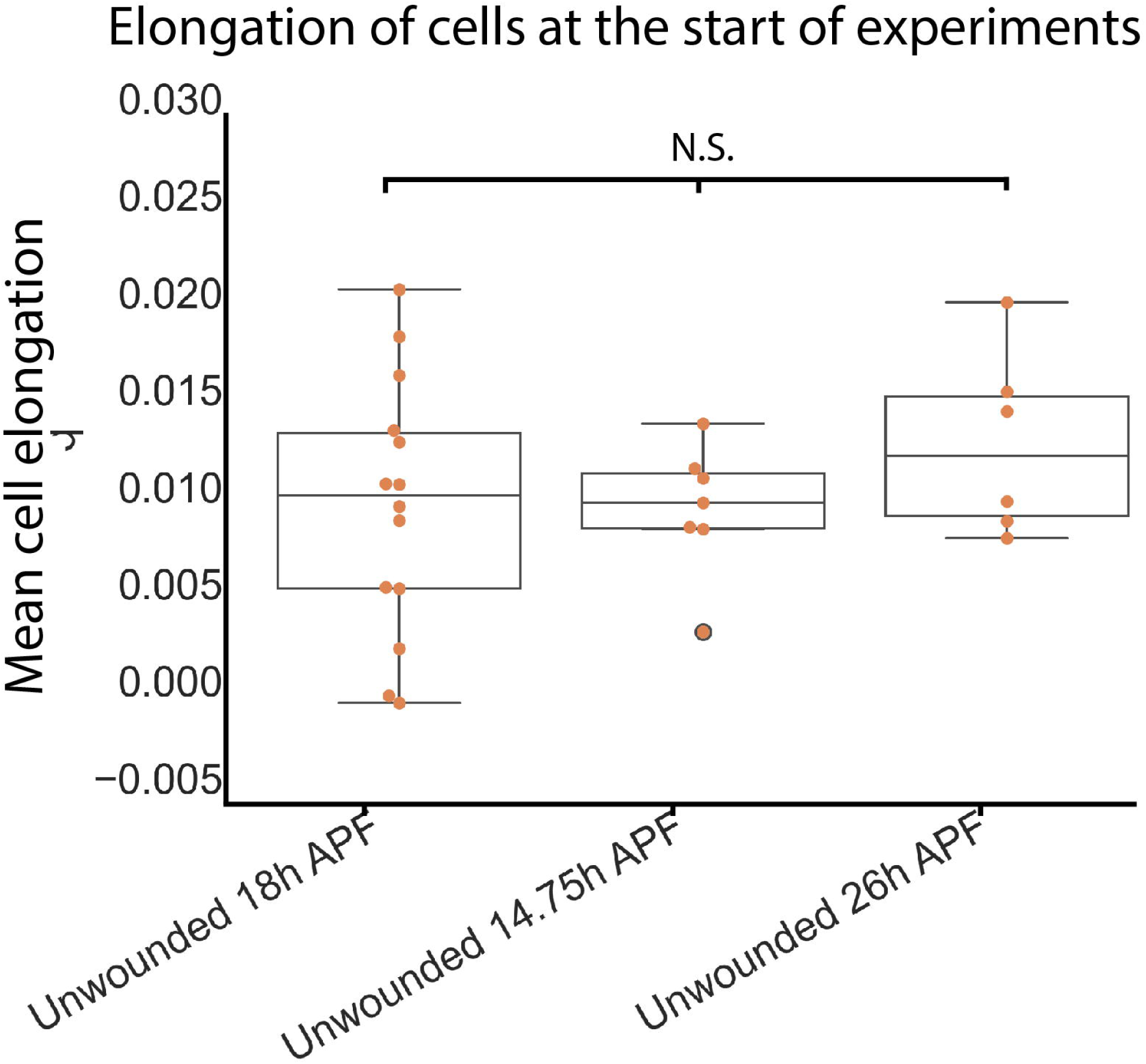
Development of pupal wings at the start of imaging, as measured via mean cell elongation to confirm consistent staging across genotypes. Box plot of the distributions of initial mean cell elongation of pupal wing tissue relative to the P/D axis of the wing. Multiple t-test were performed but without the commonly used p value corrections as we are more concerned about type II errors. (n=14 18h APF Unwounded, n=7 14.75h APF Unwounded, n=6 26h APF Unwounded)

## Supplemental Movie Legends

*Movie S1 - **Time-lapse imaging of a small wound in the pupal epithelium over 3 hours. -** Projected from a 3D stack using the stack focus algorithm with a radius of 5 pixels. Green indicates E-cadherin-GFP and magenta indicates Histone2-RFP. Scale bar: 20μm.*

*Movie S2 - **Time-lapse imaging of a large wound in the pupal epithelium over 3 hours. -** Projected from a 3D stack using the stack focus algorithm with a radius of 5 pixels. Green indicates E-cadherin-GFP and magenta indicates Histone2-RFP. Scale bar: 20μm.*

*Movie S3 - **Time-lapse imaging of a large Trpm** ^RNAi^ **wound in the pupal epithelium over 3 hours. -** Projected from a 3D stack using the stack focus algorithm with a radius of 5 pixels. Green indicates E-cadherin-GFP and magenta indicates Histone2-RFP. Scale bar: 20μm.*

*Movie S4 - **Time-lapse imaging of a large JNK knockdown wound in the pupal epithelium over 3 hours. -** Projected from a 3D stack using the stack focus algorithm with a radius of 5 pixels. Green indicates E-cadherin-GFP and magenta indicates Histone2-RFP. Scale bar: 20μm.*

*Movie S5 - **Time-lapse imaging of a large rpr wound in the pupal epithelium over 3 hours. -** Projected from a 3D stack using the stack focus algorithm with a radius of 5 pixels. Green indicates E-cadherin-GFP and magenta indicates Histone2-RFP. Scale bar: 20μm.*

## Materials and Methods

### Drosophila Stocks and Husbandry

*Drosophila* stocks were raised and maintained on Iberian food according to standard protocols (Greenspan, 1997). All crosses were performed at 25°C. The following *Drosophila* stocks were used: *ubiquitous-E-cadherin-GFP* (B#60584), *ubiquitous-Histone2-RFP* (B#23651), *Act-Gal4* (B#4414), *Tub-Gal80^ts^* (B#7108), *UAS-Trpm^RNAi^* (B#31672), TRE-DsRed (B#59011), *UAS-bsk^DN^* (B#9311), *UAS-GCamp7f* (gift from John Gillespie, University of Bristol, UK), *Srp-Gal4* (Brückner et al., 2004), *UAS-rpr* (B#5824) and UAS-nuclear-red-stinger ((a gift from Brian Stramer, King’s College London (Barolo et al., 2004)). *Drosophila* mutants and transgenic lines were obtained from the Bloomington Stock Centre unless otherwise stated.

### Confocal Imaging and wounding

*Drosophila* pupae were aged to 18h APF at 25°C unless stated otherwise. Dissection, imaging, and wounding were all performed as previously described (Weavers et al., 2018). The time-lapse movies were generated using a SP8 Leica confocal. Each z-stack slice consisted of a 123.26 x 123.26μm image (512 x 512 pixels) with a slice taken every 0.75μm. The flies were wounded on a wide-field microscope which has a nitrogen-pumped micropoint ablation laser (tuned to 435 nm, Andor) attached before being quickly transferred to the confocal microscope (Turley et al., 2023)

For the genetic perturbation experiments we use the Gal4-UAS system to explore the role of these genes in wound healing (McGuire et al., 2004). As many of these genetic manipulations are key to development and are lethal or partially lethal, we used the temperature sensitive *Tub-Gal80^ts^* to control expression of the UAS driven gene (McGuire et al., 2004). This requires us to maintain the genetically perturbed flies at 18°C during their development before shifting them to 29°C prior to imaging. For fly lines where the calcium wave or JNK signalling were to be blocked, we shifted flies to 29°C once they had developed to pre-pupae for 14.75 hours APF such that they had developed to the same stage as pupae aged at 18h APF at 25°C. For immune ablation fly lines this was still lethal therefore they were maintained at 18°C for 21hours then shifted to 29°C for 5 hours which ablated the immune cells, and the pupae developed to the same stage used in previous experiments. We compare the results of these genetically perturbed pupae to control flies which had undergone the same temperature shift schedule.

We confirmed the effectiveness of our genetic perturbations in various ways: to observe the calcium wave, we imaged a single plane in the epithelium of flies expressing UASGCamp7f (Weavers et al., 2019) with a rapid frame rate of 0.651ms/frame to catch this flash and spreading wave for up to a minute post wounding. Loss of JNK signalling in *UAS-bsk^DN^*flies was demonstrated using the downstream reporter TRE-DsRed. A z-stack of the wound was taken 5 hours post wounding for both control and knockdown tissues. The effectiveness of our immune cell ablation was seen using flies expressing nuclear-red-stinger in hemocytes, with a z-stack taken 45 mins after wounding.

### Image analysis

Image analysis was performed using imageJ and python. The code used to quantify cell behaviours and make figures for paper can be found at our GitHub repository https://github.com/turleyjm/woundHealing (databases.py, paper_BiologyWound.py). Upon publication, videos and dataset used, will also be publicly available and made available upon email request. Further details on the deep learning algorithms used to extract cell behaviours can be found in our previous publication (Turley et al., 2023). A description of the methods used is given below.

### Wound, migration, shape changes and division density measurements

We developed our own pipeline to analyse the 3 key cell behaviours involved in wound closure. Many of the main aspects of this data processing have been discuss in a previous publication (Olenik et al., 2023; Turley et al., 2023). This workflow comprises a series of automated deep, learning, and other algorithms to extract out this information with minimal human input. We quantify 4 properties from the movies wound area, cell velocity, cell shape and cell divisions. First a deep learning model called U-NetTissue segments the tissue into tissue/non-tissue binary masks. Non-tissue could be either a wound or parts of the tissue that are above or below our frame of reference. Sometimes minor hand corrections are needed to make the masks more accurate mostly when wounds have almost closed, and images become very noisy due to wound debris and immune cells. These masks of wounds closing can be used to not only measure wound area over time but also used as our frame of reference to measure cell behaviours contributing to healing.

We measure the cell migration via a single particle tracking algorithm called TrackMate. This is deployed on the 3D stack of cells nuclei (via the Histone2-RFP channel). By tracking these nuclei, we can quantify cell migration. The mean velocity is calculated and minused off the individual velocities for each nucleus to remove the general migration of the tissue and only quantify the deviations from this flow. Using the wound mask discussed above we can find the centre point of the wound and then calculate the radial component of the deviation velocity of cells in the tissue relative to the central point. This is done for every cell at every timepoint.

Next, we will calculate the elongation and orientation of each cell. This is done by focusing the 3D stack converting it into a 2D image which will be inputted to a deep learning model called U-NetBoundary, this detects cell boundaries. To maximise information input to our network we us our own focusing algorithm which is modified from similar stack focusers (Turley et al., 2023). Our method returns a RGB image with the most in focus pixels in green, then above and below pixels in red and blue respectively. Once the boundaries have been detected a watershed algorithm is applied via Tissue Analyzer (Etournay et al., 2016; Olenik et al., 2023; Turley et al., 2023). From the cell boundaries we calculate the q-tensor for each cell. This is a dimensionless second rank tensor which contains information above the elongation and orientation of each cell (Olenik et al., 2023). We compute the q-tensor which is defined as the mean elongation of cells towards the wound. Below is the derivation of this property:

For each cell *i* we calculate its q-tensor

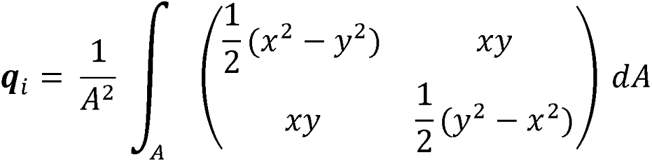

Where *A* is the area of the cell and *dA* = *dxdy*. ***q*** can be rewritten as

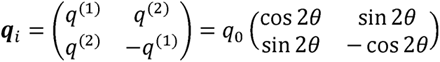

Where *θ* is the orientation of the shape. We define ⟨**Q**⟩ as the mean q-tensor over the tissue and we can calculate deviations from the mean for each cell δ**q***_i_* = **q***_i_* – ⟨**Q**⟩ removing the nematic order from the tissue. Next, we can rotate these deviation q-tensors such that the axis they are measured along aligns with radial axis from centre of the wound. Then, we take an average of them which gives δ***Q***and the component δ*Q*^(1)^ can be extract from this tensor and is the mean cell elongation relative to the wound. If δ*Q*^(1)^ > 0 then cells are on average elongated towards the wound and δ*Q*^(1)^ < 0 if perpendicular.

Lastly, we detect cell divisions using are final deep learning algorithm U-NetDivision we have recently published a paper focusing on this model so further details can be found their (Turley et al., 2023). The network can accurately find cell divisions from dynamic 2-channel (*Ecadherin-GFP* and *Histone2-RFP*) videos consisting of 5 frames. The AI algorithm finds both the position and time that the divisions occur.

Now we have extracted information about the wound and 3 cell behaviours from the videos we can now quantify their properties in space and time relative the wound. First, to calculate division density relative the wound we then calculated the distance from the edge of a wound to the divisions using a distance transform (Fabbri et al., 2008). Now we can find all the divisions in a band of a given radius and width. To quantify the density of divisions, we divide the number of divisions by the area of the band. Using both the distance transform and our tissue mask, we can work out the area of the tissue that is in each band. Once the wound has closed, we can no longer perform a distance transform using the wound edge, so we instead take the centre of the last timepoint before the wound closes. This point is the wound site and is where we take our distance transform from. As the tissue is still developing and moving, we track this point over time using the tracks the software TrackMate quantified (Tinevez et al., 2017). By calculating the average velocity of the cells around the wound site, we can track this point and use this as our frame of reference to measure the distance from the wound site.

For quantifying cell migration and shape changes we use the distance transform to find all the cells in a band of a given radius and width. Then, take the mean of either their radial component of the deviation in velocity of cells or cell elongation relative to the wound (δ*Q*^(1)^). As the q-tensor is a dimensionless quantity for easy of comparison we normalised the maximum absolute value of δ*Q*^(1)^ for large wounds.

## Acknowledgements

We would like to thank members of the Weavers, Martin, Chenchiah and Liverpool groups for helpful discussion. We also thank the Wolfson Bioimaging Facility (Bristol, UK), particularly Stephen Cross, for help setting up pyimagej, for helpful conversations and sharing useful resources. We are grateful to the Drosophila research community, Flybase and the Bloomington Stock Centre (Indiana, US), for various fly lines/reagents. JT, IC and TBL would like to thank the Isaac Newton Institute for Mathematical Sciences, Cambridge, for support and hospitality during the programme New statistical physics in living matter: non equilibrium states under adaptive control, where some of the work on this paper was undertaken. This work was supported by EPSRC grant EP/R014604/1. This research was funded by the MRC GW4 DTP PhD programme (scholarship to JT), Eric and Wendy Schmidt AI in Science Postdoctoral Fellow to JT, a Wellcome Trust and Royal Society Sir Henry Dale Fellowship to HW and a Wellcome Trust Investigator Award to PM.

